# Structure of the *Legionella* Dot/Icm type IV secretion system *in situ* by electron cryotomography

**DOI:** 10.1101/085977

**Authors:** Debnath Ghosal, Yi-Wei Chang, Kwangcheol C. Jeong, Joseph P. Vogel, Grant J. Jensen

## Abstract

Type IV secretion systems (T4SSs) are large macromolecular machines that translocate protein and DNA and are involved in the pathogenesis of multiple human diseases. Here, using electron cryotomography (ECT), we report the *in situ* structure of the Dot/Icm type IVB secretion system (T4BSS) utilized by the human pathogen *Legionella pneumophila*. This is the first structure of a type IVB secretion system, and also the first structure of any T4SS *in situ*. While the Dot/Icm system shares almost no sequence homology with type IVA secretion systems (T4ASSs), its overall structure shows remarkable similarities to two previously imaged T4ASSs, suggesting shared aspects of mechanism. However, compared to one of these, the negative-stain reconstruction of the purified T4ASS from the R388 plasmid, it is approximately twice as long and wide and exhibits several additional large densities, reflecting type-specific elaborations and potentially better structural preservation *in situ*.

## Introduction

Type IV secretion systems (T4SSs) are found ubiquitously in Gram-negative and Gram-positive bacteria as well as in some archaea (Wallden et al, 2010). They exchange genetic material within and across kingdoms and translocate virulence factors into host cells (Chandran Darbari & Waksman, 2015). T4SSs have been classified into two major groups, type IVA and type IVB (Chandran Darbari & Waksman, 2015; Sexton & Vogel, 2002). Representative examples of T4ASSs include many conjugative plasmids such as F, RP4, R388, and pKM101 and the VirB T4SS (VirB1-11 and VirD4) of the plant pathogen *Agrobacterium tumefaciens* (Chandran Darbari & Waksman, 2015; Christie et al, 2005). The VirB system is one of the best characterized T4ASSs and consists of a lytic transglycosylase (VirB1), pilins (VirB2 and VirB5), inner membrane proteins (VirB3, VirB6, VirB8), ATPases (VirB4, VirB11 and VirD4) and three factors (VirB7, VirB9, VirB10) that span the inner and outer membrane (Chandran Darbari & Waksman, 2015).

Based on major differences in composition and sequence, a subset of T4SSs were designated as T4BSSs (Sexton & Vogel, 2002). T4BSSs include the IncI conjugative plasmids R64 and ColIb-P9 and the Dot/Icm (defective in organelle <u>t</u>rafficking/<u>i</u>ntra<u>c</u>ellular <u>m</u>ultiplication) system of the pathogens *Legionella pneumophila, Coxiella burnetii*, and *Rickettsiella grylli* (Chandran Darbari & Waksman, 2015; Christie et al, 2005; Segal et al, 2005). In the case of *L. pneumophila*, the Dot/Icm system translocates more than 300 effector proteins into host cells (Isaac & Isberg, 2014), thereby allowing the pathogen to survive and replicate within phagocytic host cells (Segal et al, 1998; Vogel et al, 1998). The Dot/Icm T4BSS is more complex than most T4ASSs as it has ~27 components versus 12. The only clear sequence homology between T4A and T4B components is the C-terminus of DotG, which matches part of VirB10 (Nagai & Kubori, 2011). The Dot ATPases (DotB, DotL, DotO) are also of the same general classes of proteins as the *A. tumefaciens* ATPases (VirB11, VirD4, VirB4). Based on relationships between ATPases, T4SSs have recently been reclassified into eight classes, with the IncI class being one of the most distinct (Guglielmini et al, 2014). How similar the structures and functions of different T4SSs are remains unclear.

Great efforts have been invested into structurally characterizing different T4ASSs using an impressive array of biochemistry, crystallography and EM (Chandran Darbari & Waksman, 2015; Chandran et al, 2009; Fronzes et al, 2009; Low et al, 2014; Pena et al, 2012; Rivera-Calzada et al, 2013). The most notable achievements include a crystal structure of parts of VirB7, VirB9, and VirB10 from pKM101 (3JQO) (Chandran et al, 2009), two cryo-EM structures of the same complex (Fronzes et al, 2009), and a negative stained EM reconstruction of the recombinantly purified VirB_3-10_ complex of the related R388 plasmid (Low et al, 2014). The features of the VirB_3-10_ reconstruction were described as consisting of a periplasmic complex (cap, outer-layer, inner-layer), linked by a relatively thin stalk to an inner membrane complex (upper tier, middle tier, and a lower tier), with the latter forming two barrel-shaped densities that correspond to the VirB4 ATPase extending into the cytoplasm. However, to date no structure has been reported for any T4SS *in situ* or any of the T4BSSs. Considering their distinct genetic organization and composition, whether and how the T4A and T4B types are structurally related remains unclear.

## Results and discussion

To generate the first three-dimensional structure of a T4BSS, here we used ECT to visualize *L. pneumophila* Dot/Icm machines directly in intact, frozen-hydrated bacteria cells. In our tomograms, we observed multiple dense, cone-shaped particles in the periplasm primarily near the cell poles (Figure 1a,b; Supplementary Movie 1). These structures exhibited the characteristic shape of a “Wi-Fi” symbol comprising two distinct curved layers, the larger just below the outer membrane and the smaller in the middle of the periplasm (Figure 1c). We also observed top views of these particles, which appeared to have two concentric rings (Figure 1d). Similar rings were observed by EM imaging of portions of the Dot/Icm complex (Kubori et al, 2014). No “Wi-Fi” particles were observed in a *L. pneumophila* strain lacking the *dot/icm* genes (Figure 1E, Supplementary Figure 1).

**Figure 1.**
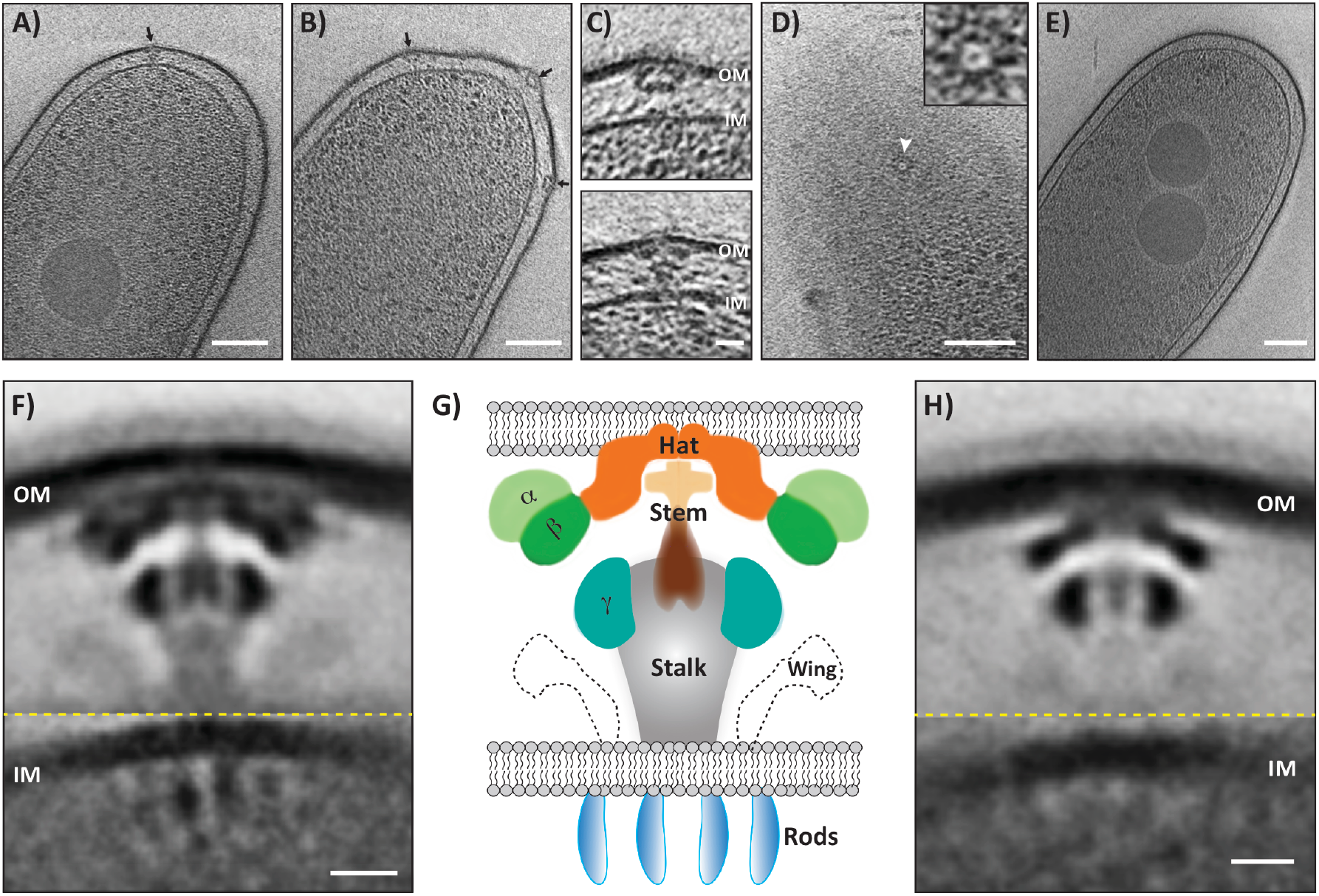
*In situ* structure of Dot/Icm T4BSS. **A,B**, Tomographic slices through intact *L. pneumophila* cells. Black arrows point to Dot/Icm particles. Scale bar 100 nm. **C**, Enlarged view of Dot/Icm particles, outer-membrane (OM) and inner membrane (IM). Scale bar 20 nm. **D**, Tomographic slices showing a top view of a Dot/Icm particle, enlarged in the inset. Scale bar 100 nm. **E**, Tomographic slice through a *L. pneumophila* cell lacking the *dot/icm* genes. Scale bar 100 nm. **F**, Subtomogram average of wild-type Dot/Icm particles. Scale bar 10 nm. **G**, Schematic representation of the subtomogram average labeling the prominent densities. **H**, Subtomogram average of a reconstituted sub-complex in the *dot/icm* deletion mutant. Scale bar 10 nm. Dotted yellow lines indicate where the outer membrane average is merged with the inner membrane average to generate the composite model.

To further investigate the molecular architecture of these complexes, we generated a subtomogram average using ~400 particles. In the initial average, substructures were resolved within the curved layers but details were lacking near the inner membrane (Supplementary Figure 2A-D). Given the previous observation of flexibility within the VirB_3-10_ complex (Low et al, 2014), we used masks to align components near the outer membrane separately from the components near the inner membrane (Supplementary Figure 2 A-D). A composite average was then constructed by juxtaposing the well-aligned regions of the outer and inner membrane averages and applying symmetry. In the final composite average, many distinct densities were resolved including a hat, alpha, and beta densities near the outer membrane; a stem, stalk, and gamma densities in the periplasmic region; and weaker densities, which we call “wings”, extending from the inner membrane into the periplasm (Figure 1F, 1G). Although of lower resolution, multiple vertical rod-like densities also appeared below the inner membrane in the cytoplasm. We estimate the local resolution of our composite model to be 2.5-4.5 nm (Supplementary Figure 3A), likely limited by inherent flexibility of the complex, as the resolution within the curves layers was the highest and the rods the lowest.

**Figure 2.**
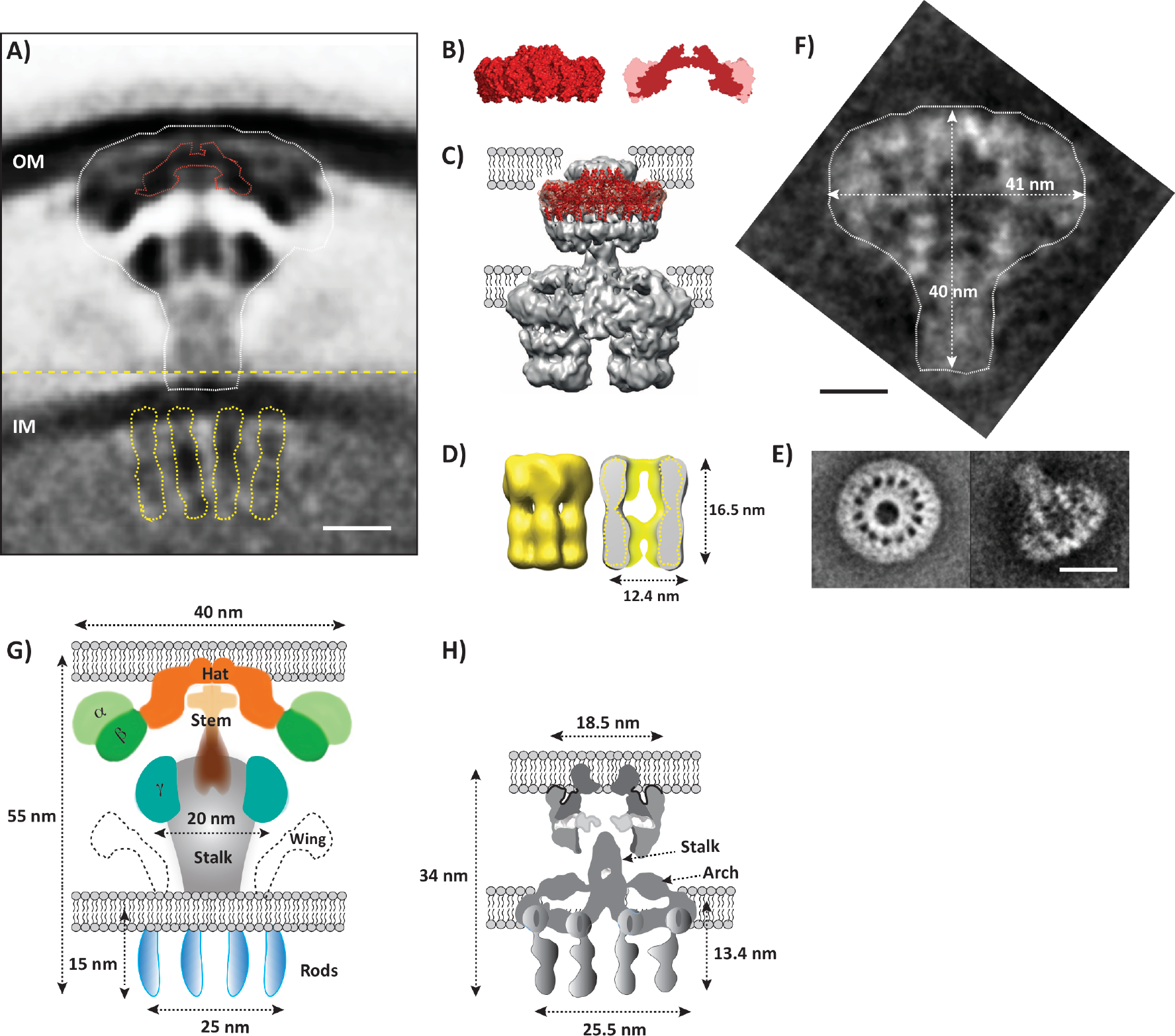
Comparison between T4ASSs and T4BSSs. **A**, Structure of the Dot/Icm complex with the outlines of existing structures of T4ASS subcomplexes superimposed. Scale bar 10 nm. **B**, Surface representation (left) and cross section (right) of crystal structure 3JQO, an outer membrane complex of parts of VirB7, 9, and 10 from the plasmid pKM101 T4ASS. Red coloured part of the cross section is density for VirB10, light-pink colour is combined density for VirB7 and 9. The outline of VirB10 density matches the hat density of the Dot/Icm structure (red dotted line in panel A, see also Supp. Movie 2). **C**, Crystal structure 3JQO fit into the VirB_3-10_ negative stain single particle reconstruction, showing its location with respect to the outer membrane (reproduced from Low et al and Chandran et al (Chandran et al, 2009; Low et al, 2014)). **D**, Isosurface of (left) and cross-section (right) through a single particle reconstruction of a purified VirB4 ATPase (EMDB accession #, EM-5505, reproduced from Pena et al (Pena et al, 2012)). Because it is a hexameric barrel-shaped structure, its cross section is two parallel rod-like densities similar to the cytoplasmic densities found in the Dot/Icm structure *in situ* (outlined in yellow in panel A). **E**, Class average images of a purified *H. pylori* T4ASS subcomplex comprising Cag3, CagM, CagT/VirB7, CagX/VirB9, CagY/VirB10 in top and side views (reproduced from Frick-Cheng et al (Frick-Cheng et al, 2016)). **F**, Same side view as in **E** but rotated and enlarged to the same scale as the Dot/Icm structure. Outline marked in white and superimposed on Dot/Icm structure in panel **A. G,H**, Schematic representations of a T4BSS (left) and a T4ASS (right, adapted from Low et al (Low et al, 2014)) showing dimensions, underlying structural similarities and differences. Scale bar for all panels except **E** 10 nm. Scale bar for panel **E**, 25 nm.

To confirm the “Wi-Fi” particles were the Dot/Icm system, we imaged a strain expressing DotC, DotD, DotF, DotG, and DotH (previously defined as the “core complex”(Vincent et al, 2006)) in an otherwise *dot/icm* null mutant strain (Figure 1H). Western blot analysis showed all five proteins were expressed at similar levels to those in the wild-type strain (Supplementary Figure 1). The subtomogram average of this reconstituted complex revealed a strong similarity to the wild-type structure as it contained the hat, beta, and gamma densities and some of the stem, but there were also major densities missing (Figure 1H and Supplementary Figure 3B, 3C). Since the “Wi-Fi” particles were not observed in a strain lacking the *dot/icm* genes, and a portion but not all of the complex reappeared upon reintroduction of the five core Dot proteins, we are confident that these particles are the Dot/Icm system rather than some other membrane complex such as the *L. pneumophila* T2SS or a different T4SS.

In T4ASSs, a “core complex” has been described consisting of three proteins with major domains in the periplasm: the inner membrane protein VirB10, the outer membrane protein VirB9, and a lipoprotein VirB7, which plays a role in the insertion of VirB9 (Chandran Darbari & Waksman, 2015). In the *Legionella* T4BSS, DotF and DotG are inner-membrane proteins, DotH is an outer membrane, and there are two lipoproteins, DotC and DotD, that function to insert DotH (Vincent et al, 2006) Markedly, and as mentioned above, among these proteins there is only one domain shared between the Dot and VirB systems: the C-terminus of DotG has clear sequence homology to VirB10 (Chandran Darbari & Waksman, 2015). Despite this paucity of homology between components, the *in situ* structure of the Dot/Icm T4BSS and the negative-stain reconstruction of the VirB_3-10_ T4ASS complex clearly share key features. First, the size and shape of the hat density in the Dot/Icm apparatus matches the VirB10 density from the crystal structure 3JQO, which contains parts of VirB7, VirB9, and VirB10 (Figure 2A-C, Supplementary Movie 2). This makes sense because the domain of VirB10 present in the crystal structure is the one with sequence homology to DotG (Supplementary Figure 4B). Thus it is not surprising that there would be a similar-shaped feature in the equivalent location of the Dot/Icm structure (as seen in the hat). Both the Dot/Icm and VirB_3–10_ structures also contain flexible stalks between the outer membrane and inner membrane complexes. Finally, the four rod-like densities in the Dot/Icm structure correspond well in size, shape and position (with respect to both the inner membrane and stalk) to the walls of the two barrels seen in the VirB_3–10_ complex, leading us to conclude that there are two similar barrels present in the Dot/Icm complex, even though they are poorly resolved (Figure 2A, 2D, 2G-H). Thus the basic architecture of the Dot/Icm system is strikingly similar to that of the VirB_3-10_ complex: each contains a hat, stalk, and two off-axis cytoplasmic barrels.

However, there are also major differences. First, the Dot/Icm complex is strikingly larger than the VirB_3-10_ complex, as it is approximately twice as wide and long (Figure 2G-H). Second, there are no densities in the VirB_3-10_ complex peripheral to the hat which might correspond to alpha and beta. Third, it is not clear if the Dot/Icm gamma density is part of what was described as the inner layer of the VirB_3-10_ complex. Fourth, the Dot/Icm structure has periplasmic wings instead of membrane-associated arches. While some of these differences are likely due to the additional factors present in the Dot/Icm system, others may reflect the loss or collapse of components in the VirB_3-10_ complex upon purification and drying. The arches of the VirB_3-10_ complex, for instance, may correspond to collapsed Dot/Icm wings, and the shorter and thinner stalk in the VirB_3-10_ complex may also be a result of collapse (the distance between the outer and inner membranes in our cryotomograms of different species of intact bacterial cells is typically ~40 nm (Chang et al, 2016; Chen et al, 2011), twice as far as in the VirB_3-10_ complex structure).

Recently, two-dimensional class average images of a negatively-stained *H. pylori* T4ASS comprising Cag3, CagM, CagT/VirB7, CagX/VirB9 and CagY/VirB10 were reported (Figure 2E-F) (Frick-Cheng et al, 2016). While the *Helicobacter* T4ASS consists of approximately the same number of components as the *Legionella* Dot/Icm T4BSS, the additional factors share no homology (Frick-Cheng et al, 2016). Despite also being purified, dried, and negatively-stained like the R388 plasmid VirB_3-10_ complex, the *Helicobacter* structure has almost exactly the same size and overall shape as the periplasmic region of the *Legionella* structure, with a large bulbous structure at one end and a stalk at the other (Figure 2A, 2E-F). Because the *Helicobacter* images were of a purified subcomplex, it was impossible at the time to assign an orientation of the structure relative to the envelope (Frick-Cheng et al, 2016). Based on the similarities to our *in situ* structure, we can now predict that the concave surface faces the outer membrane and the stalk points in the direction of the inner membrane.

In summary, we have revealed the first *in situ* structure of a T4SS and shown that despite very little sequence homology between representative T4ASSs and T4BSSs (Supplementary Figure 4A,B), their basic architectures and therefore likely secretion mechanisms are remarkably similar. They are much more similar than different when compared to the structure of other secretion systems. Type III secretion systems, for example, consist of a series of rings in the inner membrane, periplasm and outer membrane connected by a central channel that serves as the conduit for protein export (Hu et al, 2015). In contrast, neither the T4ASS nor the T4BSS exhibit an obvious tube-like channel along the symmetry axis through which substrates might be transported. Although informative within its own right, the *in situ* Dot/Icm structure also sets the stage for future work identifying each protein in the complex and elucidating how this elaborate nanomachine assembles and functions.

## Materials and methods

### Strains, growth conditions and mutant generation

All experiments mentioned here were performed using the *L. pneumophila* Lp02 strain (*thyA hsdR rpsL*), which is a derivative of the clinical isolate *L. pneumophila* Philadelphia-1. *L. pneumophila* strains were grown on ACES [N-(2-acetamido)-2-aminoethanesulfonic acid]-buffered charcoal yeast extract agar (CYE) or in ACES-buffered yeast extract broth (AYE) supplemented with ferric nitrate and cysteine hydrochloride. Since Lp02 is a thymidine auxotroph, cells were always grown in the presence of thymidine (100 μg/ml). JV5443 is a derivative of Lp02 lacking the *dot/icm* genes (JV5319) that was transformed with plasmid pJB4027, which expresses *dotD, dotC, dotH, dotG*, and *dotF*.

### Sample preparation for electron cryotomography

*L. pneumophila* (Lp02) cells were harvested at early stationary phase (OD600 of ~3.0), mixed with 10-nm colloidal gold beads (Sigma-Aldrich, St. Louis, MO) precoated with BSA, and applied onto freshly glow-discharged copper R2/2 200 Quantifoil holey carbon grids (Quantifoil Micro Tools GmbH, Jena, Germany). Grids were then blotted and plunge-frozen in a liquid ethane/propane mixture (Chang et al, 2016) using an FEI Vitrobot Mark IV (FEI Company, Hillsboro, OR) and stored in liquid nitrogen for subsequent imaging.

### Electron tomography and subtomogram averaging

Tilt-series were recorded of frozen *L. pneumophila* (Lp02) cells in an FEI Titan Krios 300 kV field emission gun transmission electron microscope (FEI Company, Hillsboro, OR) equipped with a Gatan imaging filter (Gatan, Pleasanton, CA) and a K2 Summit direct detector in counting mode (Gatan, Pleasanton, CA) using the UCSF Tomography software (Zheng et al, 2007) and a total dose of ~100 e/A^2^ per tilt-series and target defocus of ~6 *μ*m underfocus. Images were aligned, CTF corrected, and reconstructed using IMOD(Kremer et al, 1996). SIRT reconstructions were produced using TOMO3D(Agulleiro & Fernandez, 2015) and subtomogram averaging was performed using PEET (Nicastro et al, 2006). Finally, the local resolution was calculated by ResMap (Kucukelbir et al, 2014). As the Dot/Icm subtomogram average exhibited at least two-fold symmetry around the central mid-line in the periplasm, we applied two-fold symmetry in those regions to produce the 2-D figures shown, but no symmetry was applied to the cytoplasmic densities due to their poor resolution.

